# Word and sentence embedding tools to measure semantic similarity of Gene Ontology terms by their definitions

**DOI:** 10.1101/103648

**Authors:** Dat Duong, Wasi Uddin Ahmad, Eleazar Eskin, Kai-Wei Chang, Jingyi Jessica Li

## Abstract

The Gene Ontology (GO) database contains GO terms that describe biological functions of genes. Previous methods for comparing GO terms have relied on the fact that GO terms are organized into a tree structure. Under this paradigm, the locations of two GO terms in the tree dictate their similarity score. In this paper, we introduce two new solutions for this problem, by focusing instead on the definitions of the GO terms. We apply neural network based techniques from the natural language processing (NLP) domain. The first method does not rely on the GO tree, whereas the second indirectly depends on the GO tree. In our first approach, we compare two GO definitions by treating them as two unordered sets of words. The word similarity is estimated by a word embedding model that maps words into an N-dimensional space. In our second approach, we account for the word-ordering within a sentence. We use a sentence encoder to embed GO definitions into vectors and estimate how likely one definition entails another. We validate our methods in two ways. In the first experiment, we test the model’s ability to differentiate a true protein-protein network from a randomly generated network. In the second experiment, we test the model in identifying orthologs from randomly-matched genes in human, mouse, and fly. In both experiments, a hybrid of NLP and GO-tree based method achieves the best classification accuracy.

## 1 Introduction

The Gene Ontology (GO) project founded in 1998 is a collaborative effort that has been providing consistent descriptions of genes and proteins across different data sources and species^[6]^. The GO database is similar to a dictionary; it contains terms referred to as GO terms. Each GO term has a definition describing some biological event.

The GO database is divided into three categories: cellular components (CC), molecular functions (MF) and biological processes (BP). The CC ontology contains terms describing the components of the cell and can be used to locate a protein. The MF category contains terms describing chemical reactions such as *catalytic activity* or *receptor binding.* These terms do not specify the genes or proteins involved in the reactions or the locations of the events. The BP category contains terms describing a series of biological events. For example, the BP term GO:0006874 has the definition “Any process involved in the maintenance of an internal steady state of calcium ions at the level of a cell.” In each category, the GO terms are organized into a tree where there is only one *root* node^[6]^. In this tree of GO terms (or GO tree), a more generic term (i.e. lyase activity) is closer to the root, whereas a more specific term (i.e. carboxy-lyase activity) is closer to a leaf node.

Because there are three GO categories in the database, there are three GO trees. Interestingly, there are edges connecting terms in different GO trees (Figure 1). The GO database can be represented as three connected GO trees.

**Figure 1:**
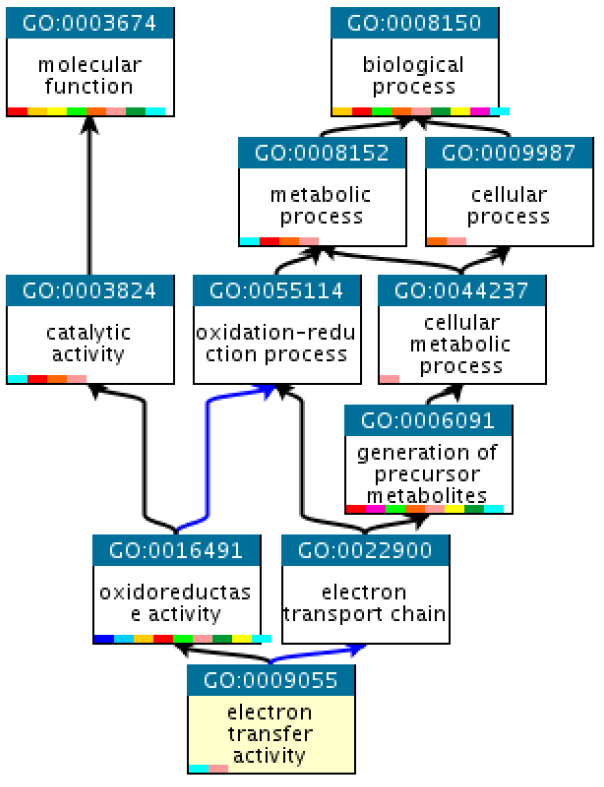
Terms shown are ancestors of GO:0009055 (yellow). GO:0003674 and GO:0008150 are root nodes for the MF and BP trees, respectively. Colors denote *part of* (blue) and *is a* (black) relationship. Snapshot is downloaded from ebi.ac.uk/QuickGO.

One application of the GO database is the comparison of two genes by first comparing the similarity of the GO terms that annotate them^[6]^. To this end, we need a good metric for comparing GO terms. To solve this problem, we need to focus on the GO trees and the definitions of GO terms. Because of the GO trees, GO terms with a direct ancestor (i.e. sibling nodes) are deemed to be more related than GO terms with a distal ancestor. Moreover, because of this design, existing methods to measure the similarity of two GO terms mostly rely on the GO trees^[16]^. Very few studies have yet to directly compare the definitions of GO terms^[19]^.

In this paper, we introduce two new solutions to measure the semantic similarity of two GO terms, by focusing on the definitions of the GO terms. Our approach is most similar to Pesaranghader <u>et al.</u>^[19]^; however, we apply neural network based techniques from the natural language processing (NLP) domain.

First, we compare words by converting them into word embeddings. We train the Word2vec model using open access articles on PubMed, so that we can represent a word as an N-dimensional vector^[17]^. Cosine similarity is used to compare two words. To compare two GO terms, we treat their definitions as two unordered sets of words, and use the weighted Modified Hausdorff Distance to measure the distance between two sets^[4]^. We name this metric w2vGO. w2vGO is entirely independent of the GO trees.

Second, we consider the word-ordering within the GO definitions. We note that *entailment* relationships exist in the GO tree (i.e. two GO terms are linked by a directed edge) (Figure 1). We train the sentence encoder InferSent^1^ using the definitions of child-parent and randomly-matched GO terms^[2]^. InferSent embeds sentences into an N-dimensional vector space and computes the probability that one sentence entails another. We name this approach InferSentGO. InferSentGO needs the GO trees for the training phase. Once the model is trained, only the definitions of GO terms are required for calculating their semantic similarities.

We compare W2vGO and InferSentGO against simDEF by Pesaranghader <u>et al.</u>^[19]^, and the following tree-based methods: Resnik, GraSM, Aggregate Information Content (AIC)^[3,23,26]^. For Resnik and GraSM, we include the random walk contribution (RWC) to their scores by using the software GOssTo^[1,29]^. We will discuss these competing methods in the Method and Appendix section. We further consider an ensemble approach AicInferSentGO by averaging the AIC and InferSentGO scores.

To compare these metrics, we conduct two experiments. In the first experiment, we test these metrics in differentiating the true protein-protein interaction network from a random network for both human and yeast. In the second experiment, we test the metrics in identifying orthologs against randomly-matched gene pairs for human, mouse, and fly. Our results show that the hybrid AicInferSentGO attains the best classification in terms of the area under the receiver operating characteristic (ROC) curve. Our software, data, and results are available at our GitHub^2^.

## 2 Method

### 2.1 Methods to measure similarity between two GO terms

Broadly speaking, existing tree-based methods are divided into two types: node-based or edge-based^[15]^. The key focus of node-based methods is the evaluation of the information content of the common ancestors for two GO terms. In brief, the information content of a GO term measures the usefulness of the term by evaluating how often the term is used to annotate a gene. Terms that are used sparingly have high information content because they are specific at distinguishing genes. Node-based methods have been shown to work well^[14,15,26]^. However, we show in section 2.4 that we can improve the accuracy by including the semantic similarities of the GO definitions.

In this paper, we choose the node-based methods Resnik and AIC as the baselines. Resnik is a classical approach for quantifying the similarity between two GO terms^[23]^. Despite being simple, Resnik has been shown to outperform some of its extensions on several test datasets^[15,21]^. AIC is recent. In their paper, Song <u>et al.</u>^[26]^ showed that AIC outperforms popular node-based methods by Jiang and Conrath^[9]^, Lin <u>et al.</u>^[13]^, Resnik^[23]^, and Wang <u>et al.</u>^[28]^.

Unlike node-based methods, edge-based approaches analyze the paths between two GO terms. For example, these approaches measures the distance (or the average distance when more than one path exists). We include the edge-based GraSM as the baseline^[3]^. We also apply Yang <u>et al.</u>^[29]^ approach (via GOssTo software) to add extra topological information from the GO tree for both GraSM and Resnik. Yang <u>et al.</u>^[29]^ referred to this extra information as the random walk contribution (RWC). In the appendix, we describe Resnik, AIC, GraSM, and RWC in detail.

Recently, Pesaranghader <u>et al.</u>^[19]^ introduced simDEF, a text mining approach to compare the definitions of two GO terms. Their work is most similar to ours; however, they do not use neural network techniques. We briefly describe the five steps in simDEF. First, simDEF counts the co-occurrences for words in the MEDLINE corpus to construct a symmetric first-order co-occurrence matrix. Values in this matrix represent how many times the word in its row appears with the word in its column. Second, simDEF extends the definition of a GO term by concatenating its definition with the definitions of its direct parents and children. Third, simDEF builds a second-order matrix from the first-order matrix^[8]^. Loosely speaking, each row in second-order matrix is an extended GO definition. Each column is the word count for a word in the definition. Fourth, simDEF applies the Pointwise Mutual Information (PMI) function on the second-order matrix^[8]^. In their paper, Pesaranghader <u>et al.</u>^[19]^ refers to this outcome as the PMI-on-second-order matrix. Here, each row represents an extended GO definition, and the entry for each row is the transformed word count for a word in the definition. Fifth, cosine distance is used to measure the similarity between two GO terms (i.e. two rows in the PMI-on-second-order matrix).

We now introduce our methods and outline their differences from simDEF.

### 2.2 Word2vec model

Here, we describe our first metric W2vGO. We use W2vGO as pure NLP technique; that is, W2vGO focuses strictly on the GO definitions and is completely independent of the GO-trees. For this reason, unlike simDEF, we do not concatenate a GO definition with the definitions of its parents and children. This extension of a GO definition requires information from the GO tree.

To compare two GO terms, W2vGO compares their definitions by treating the definitions as two unordered sets of words. To solve this problem, we want to first be able to compare two words. To this end, we use the word embedding model Word2vec^[17]^. Word2vec is a distributional linguistic model. Loosely speaking, Word2vec analyzes how often words co-occur. However, Word2vec is very different from simDEF co-occurrences matrices.

The Word2vec model converts a word into an *N*-dimensional vector^3^. These vectors are known as word embeddings. Word2vec transforms similar words into similar vectors, thus enabling us to quantify the similarity between two words by computing the Euclidean distance or cosine similarity. At the heart of the Word2vec is a neural network model with one input layer, one hidden layer, and one output layer^[17]^.

Loosely speaking, one can view the Word2vec model as a prediction problem^[24]^. First, a word *w* from the input layer is mapped into an *N*-dimensional vector at the hidden layer. Word2vec learns the values for the hidden layer based on the co-concurrences between *w* and its neighboring words. The key idea is to predict the vectors for the surrounding context words based on the vector for *w*.

The purpose of this paper is not to dissect the Word2vec model; we are interested in adopting this model to measure the similarity of GO terms and compare it with other methods. Interested readers are encouraged to read the original paper by Mikolov <u>et al.</u>^[17]^, and the introduction by Rong^[24]^.

#### 2.2.1 Measure similarity of two words using Word2vec

The training data influences the application of word embedding model. Existing pretrained Word2vec models are often made by corpora collected from news, books, or the Internet. To obtain suitable word vectors, in this work, we train the Word2vec to recognize biological words. We set the dimension *N* = 300 and use 20 GB of data from open access articles on PubMed. The raw count of unique and repeated words is 14,526,527,855. We remove words which appear less than 25 times in the whole training data, thus reducing the final number of unique words to 986,615. We keep stop-words and symbols like + and — in the data because they may have important biological meanings. We use the Python library gensim to train the Word2vec model^[22]^. A simple Python user interface is available at our GitHub.

There are two important details here. First, the training data do not contain definitions of GO terms found in the GO database. This helps us avoid data reusing. Second, theoretically speaking, Word2vec model can be trained on the GO terms in the PubMed data, so that one can convert a GO term into a vector. Unfortunately, the IDs of the GO terms are not used too often in published papers, and detecting definitions of GO terms in papers is a different type of research problem^[27]^. For these reasons, we use the Word2vec model as a metric to compare two biological words.

To compare two vector representations of two words, we use the cosine similarity, because it is bounded; whereas, Euclidean distance is not. We define the function w2v(*z, v*) as the similarity score of two words *z, v*.

#### 2.2.2 Measuring similarity of two GO terms using Word2vec

A GO term comes with a definition that is usually one or two sentences describing some biological feature. For example, GO:0003700 has the definition: “Interacting selectively and non-covalently with a specific DNA sequence in order to modulate transcription. The transcription factor may or may not also interact selectively with a protein or macromolecular complex.”

When a GO term definition has more than one sentence, we concatenate these sentences into the same sentence by ignoring the period symbol. For example, the two sentences for GO:0003700 is considered as one long sentence.

Thus, the task to compare two GO terms reduces to the problem of comparing their definitions which are two sentences. Suppose that GO terms *a, b* have sentences *Z, V* as their definitions respectively. We treat two sentences *Z* = “*z*_1_ *z*_2_ *z*_3_ … *z_N_*” and *V* = “*v*_1_ *v*_2_ *v*_3_ *… v_M_*” as two unordered sets of words *Z* = {*z*_1_*, z*_2_ *… z_N_*} and *V* = {*v*_1_*,v*_2_ *… v_M_*}.

To measure the similarity of sentences *Z* and *V* (or in other words, term *a* and *b*), we use the metric

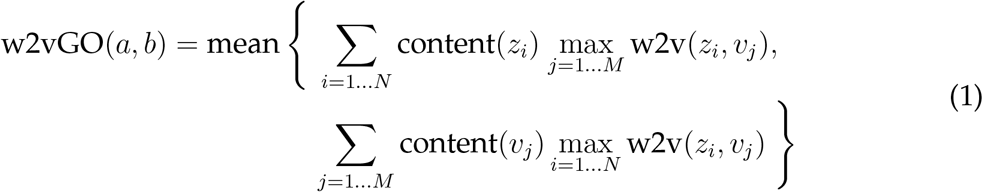

where content(*w*) is the weight of the word *w* and is often used to distinguish common words from rare ones. The weights of words can help avoid the influence of hub-words (i.e. words such as *cell, DNA, activity*) that are ubiquitously associated with many other words^[11]^. content(*w*) is very similar to the IC function^[12]^

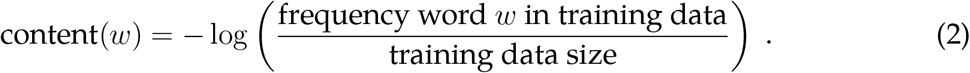

Because GO definitions are often short, we hope that the accuracy of w2vGO does not suffer too much from the removal of word-ordering. Surprisingly, this assumption holds true in several instances (Table 2). There are more sophisticated models that consider the word-ordering in the sentences; we will consider one of these methods in the next section.

In any case, we have defined a metric to measure the two GO terms *a, b* with definitions *Z, V* under the Word2vec paradigm. w2vGO(*a, b*) ranges from −1 to 1 because w2v(*z, v*) ranges from −1 to 1.

### 2.3 InferSent model

In the previous section, we have ignored the word-ordering in the sentences, treating them as sets of words. In this section, we briefly explain InferSent, a model that focuses not only on the word embeddings but also on the word-ordering in the sentences^[2]^. InferSent is thus different from simDEF which also ignores the word-ordering.

Loosely speaking, InferSent is similar in spirit to Word2vec; instead of words, InferSent identifies the relationship between two sentences. One training sample for InferSent consists of two sentences having some relationship *R*. Consider a simple classification where *R* = *entailment* or *neutral.* In this case, InferSent expect two classes of input. The first class *entailment* will have two sentences where the first *entails* the second; for example, “cat is napping on the mat” *entails* “cat is not running”. *Entailment* is a one-directional relationship, because “cat is not running” does not *entail* “cat is napping on the mat”. The second class *neutral* will have two unrelated sentences like “cat is napping on the mat” and “boy is watching the cat”. Unlike *entailment, neutral* is a bi-directional relationship.

InferSent is a classification model based on the neural network architecture; its full description is at Conneau et al.^[2]^. Here, we briefly mention the first layer of this network. InferSent’s first layer is the word vectors for words in the entire training dataset. Conneau <u>et al.</u>^[2]^ uses the GloVe word vectors^[18]^. However, to obtain the best result, these word vectors should be specific to biology. In this paper, we use the Word2vec vectors in section 2.2.1.

#### 2.3.1 Measuring similarity of two GO terms using InferSent

InferSent takes two sentences as one training sample. In this paper, the two sentences will be the definitions of two GO terms *a, b.* We will define two categories *entailment* and *neutral,* and estimate the probability 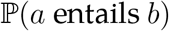. This metric allows us to gauge the semantic similarity for *a* and *b*. We choose this option because *entailment* relationship exists in the GO tree. Child-parent GO terms are linked by a one-directional relationship like “is a”, “part of”, “regulates”, “negatively regulates”, and “positively regulates”. For example, the term GO:1900237 “positive regulation of induction of conjugation with cellular fusion” *entails* the term GO:0010514 “induction of conjugation with cellular fusion.”

To prepare the *entailment* dataset, for all three ontology categories, we randomly pair each GO term with one of its parents. To ensure that these child-parent GO terms are indeed similar in meaning, we compute the median AIC score for each category and retain pairs having scores above the median. Our final dataset contains 17,226 pairs. We treat the three ontology categories as one single dataset when training the InferSent model.

To create the *neutral* dataset, we make two types of unrelated pairs. For the first type, we randomly pick about half the number of GO terms in the *entailment* dataset. For each term *c* in this set, we pair it with a randomly chosen GO term *d* in the same ontology category. For the second type, we pair the same term *d* with another randomly chosen term *e*. This sampling scheme improves the training by allowing some GO terms to be seen more than once under different circumstances.

Two unrelated GO terms should have 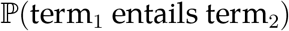 and 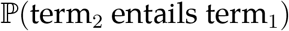 near zero. For each *neutral* pair, we create two different samples for InferSent. The first sample will be the pair (definition of term_1_, definition of term_2_), and the second sample will be the pair (definition of term_2_, definition of term_1_). Our final *neutral* dataset contains 35,044 pairs of sentences. Since the *neutral* dataset is nearly twice as large as the *entailment* dataset, when training InferSent, we weigh ratio 1:2 for the class *neutral* and *entailment.*

The code to train InferSent is available at our GitHub. We attained 96.93% accuracy in the validation set. We emphasize that InferSent relies on the GO tree only for its training phase. Once the model is trained, it requires only the GO definitions as the input for prediction.

When terms *a* and *b* are unrelated, we expect 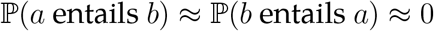. When GO term *a* is the child of term *b*, we expect 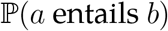 to be high, whereas 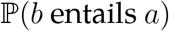 may be low. However, in this case, we still want the similarity between *a, b* to be high. For this reason, to measure the semantic similarity for two GO terms, we use the metric

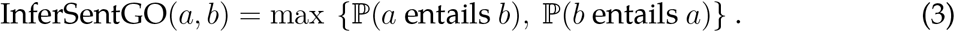

InferSentGO(*a, b*) ranges from 0 to 1.

### 2.4 Combining node-based and NLP methods

In this section, we discuss a few examples and motivate the need for combining node-based and NLP methods. Here, we choose the node-based Resnik and AIC, because GraSM and RWC do not yield good results (Table 2).

In essence, both Resnik and AIC focus on the fraction of shared ancestors and weigh this ratio by the IC values. Sometimes, even when two terms are very similar in meaning, they can have a low score. For example, consider the terms GO:0005887 and GO:0016021 in the CC ontology (Table 1 Row 1). This is a child-parent pair, having almost identical definition. However, the Resnik and AIC score are not high enough. In a sample of 3,244 child-parent pairs in the CC ontology, we found the median score to be 5.5933 and 0.9278 for Resnik and AIC, respectively.

**Table 1:**
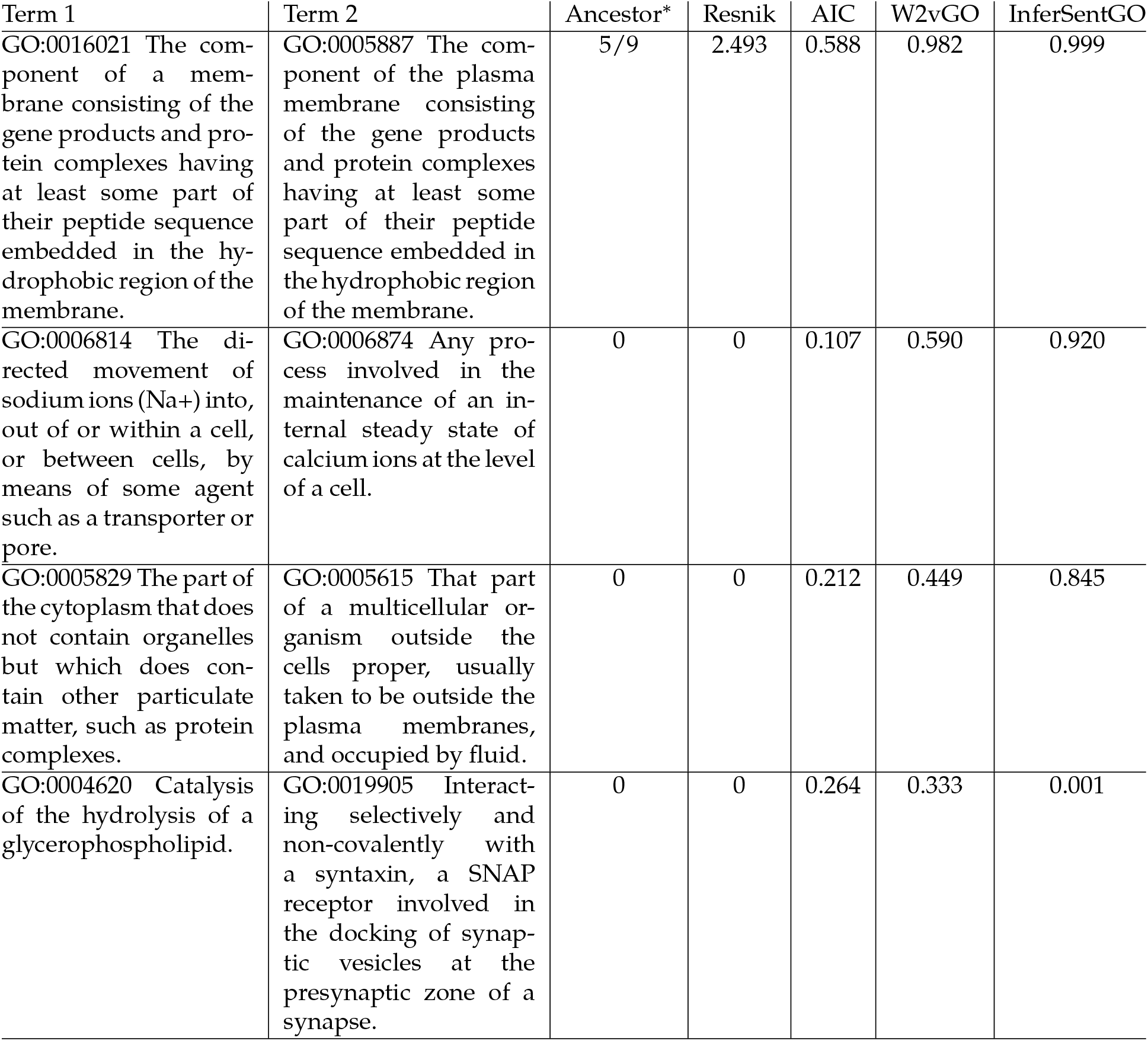
A few examples to compare GO similarity scores. * Fraction of shared ancestors, with 0 indicates terms share only the root node.

In the second example (Table 1 Row 2), GO:0006814 and GO:0006874 are unrelated, sharing only the root node. Interestingly, one can argue that both terms are not entirely distinct because they both mention the regulation of ions. Similarly, in the third example (Table 1 Row 3), GO:0005829 and GO:0005615 share only the root node, but both mention fluid containing protein complexes. In both examples, W2vGO on its own or an average of AIC and InferSentGO may give a more satisfying score. InferSentGO and AIC on their own may underestimate and overestimate the similarity, respectively. Resnik on its own is not the best because when terms share only the root node, the similarity score is the IC of the root which is 0.

In the final example (Table 1 Row 4), GO:0004620 and GO:0019905 share only the root node and truly are different in meaning. Here, Resnik and InferSentGO give more reasonable scores than AIC and W2vGO do.

These examples suggest that no one method is always the best, and that we need to combine node-based and NLP approaches. To this end, taking an average of AIC and InferSentGO is reasonable. Empirically, from the examples, we have seen that this average produces a reasonable value. Theoretically, only AIC and InferSentGO scores are in the same [0, 1] range; whereas Resnik and W2vGO range are [0, ∞] and [−1, 1]. We note that simDEF range is [−1, 1], making it incompatible with node-based Resnik and AIC.

For two GO terms *a, b*, we introduce the metric

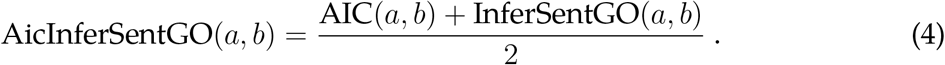

From AIC’s perspective, AicInferSentGO improves the distinction between child-parent and randomly-matched GO terms, especially in the CC and MF ontology (Figure 2). From InferSentGO’s perspective, AicInferSentGO gives a more continuous score, allowing for a better resolution when comparing terms. InferSentGO on its own tends to give a stiff 0/1 score.

**Figure 2:**
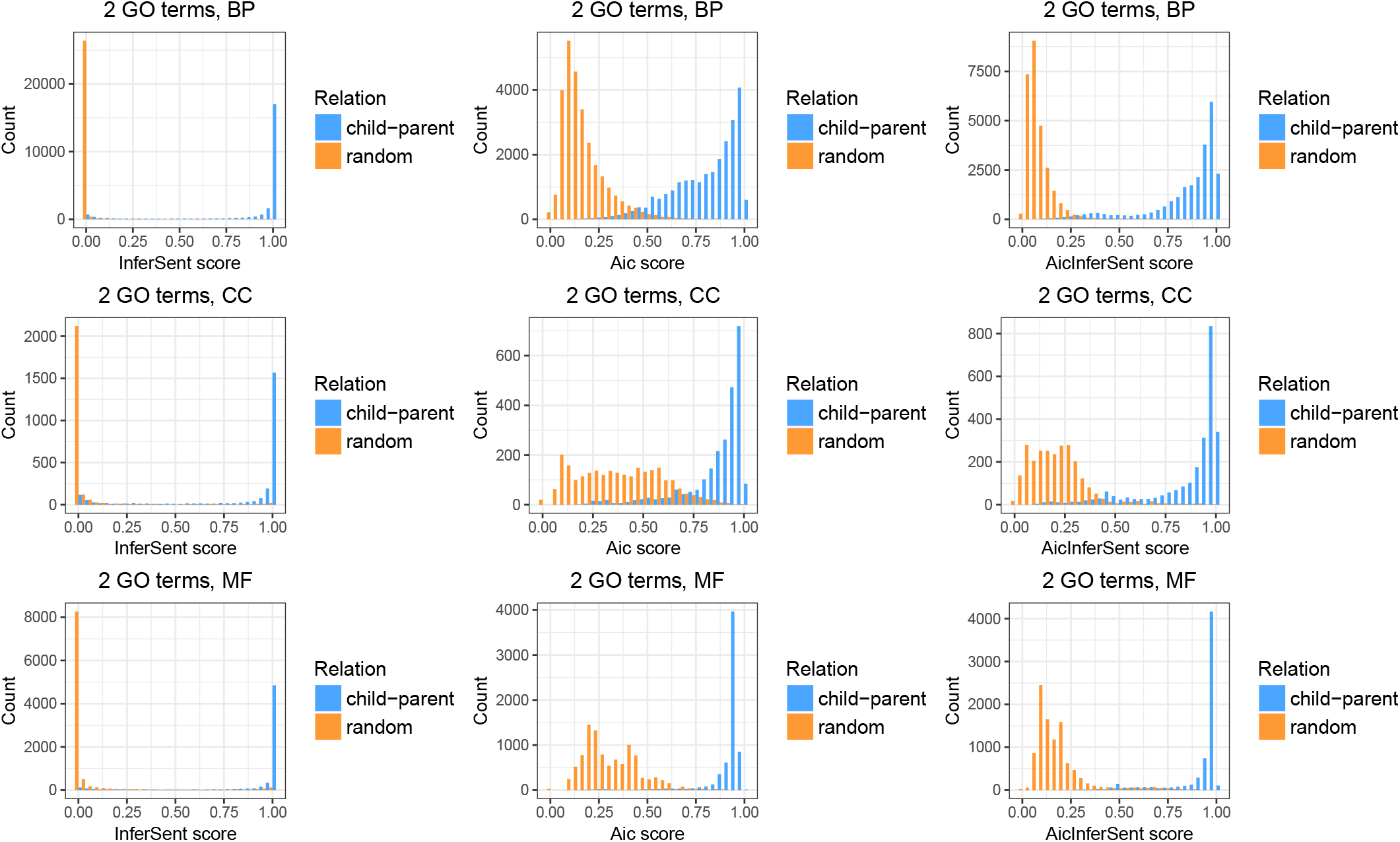
Similarity scores for 33020 child-parent GO terms, and 40419 randomly-matched GO terms. The number of pairs are 51931, 4986, and 16522 for BP, CC, and MF ontology respectively. From AIC’s perspective, the hybrid method AicInferSentGO improves the distinction between child-parent and randomly-matched GO terms. From InferSentGO’s perspective, AicInferSentGO gives a more continuous score.

### 2.5 Measuring similarity of two genes

To assess the performance of the different GO metrics, we will use them to compare genes. A gene is annotated with several GO terms from the three GO categories. For example, the gene HOXD4, which is important for morphogenesis, is annotated by these GO terms GO:0003677, GO:0003700, and GO:0006355. Thus, we can view any gene *A* as a set of GO terms. A GO term *a* is in the set *A* (i.e. *a* ∈ *A*) if *a* is used to annotate *A*.

To assess the similarity between two genes *A* and *B*, we must compare two sets of GO terms. There are many metrics for this task^[16]^. Here, we use the Modified Hausdorff Distance (MHD) and the Best Max Average distance (BMA). MHD is a traditional metric for comparing two sets, and was often used in image processing (i.e. to compare two sets of pixels)^[4]^. BMA has been shown to be better than taking the maximum or minimum of all pairwise distances for the elements in the two sets^[20]^.

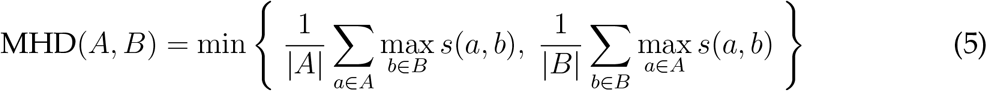

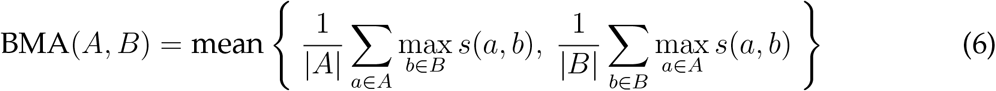

In the above, the function *s*(*a, b*) is a generic placeholder for measuring the similarity of GO terms *a* and *b*. For example, if one uses Resnik, AIC, or W2vGO metric then *s*(*a, b*) = Resnik(*a, b*), AIC(*a, b*) or w2vGO(*a, b*) respectively.

## 3 Results

We compare the GO metrics. Because genes are annotated by GO terms, good GO metrics should differentiate similar genes from unrelated genes well. Hence, we conduct two experiments. First, we test the GO metrics in identifying true protein-protein interactions. Second, we test the metrics in identifying orthologs in human, mouse, and fly. We download the GO term definitions and GO annotations from the Gene Ontology (geneontology.org). We download the orthologs from Ensembl (ensembl.org/biomart). The source code, data, and results in this section are available at our GitHub.

### 3.1 Human protein-protein interaction network

We use the 6031 protein-protein interaction (PPI) data prepared by Mazandu and Mulder^[15]^. We trim this data further, keeping only human proteins that can be mapped to some genes via UniProt (uniprot.org). Next, to avoid data reusing, we remove electronically inferred GO terms (removing terms with tag IEA, NAS, NA, NR)^[20]^. To keep only genes that are well studied, we retain only genes with at least one GO term in each ontology (BP, CC, and MF). The final data has 2593 pairs.

Like in Mazandu and Mulder^[15]^, we want to compare how well each metric differentiates a true PPI network (positive set) from a randomly made PPI network (negative set). We follow the procedure by Mazandu and Mulder^[15]^. We make the positive and negative sets to have the same number of edges. For the negative set, we randomly assign edges between proteins that do not interact in the real PPI network. The real and random PPI network have the same proteins; we only require that they have different interacting partners. For each PPI network, to compute the similarity scores of the edges (i.e. pairs of proteins), we use Eq. 5 and 6 with *s*(*a, b*) being one of the GO metrics in Table 2.

**Table 2:** AUCs for classifying human protein-protein interactions. Bold font indicates the best value in each column. *Joint analysis: When comparing genes, we keep their entire GO annotations, effectively treating the BP, CC, MF ontologies as connected GO trees. +Pvalue of the Hanley-McNeil test to compare AUCs of other methods against BMA+AicInferSentGO.

| Set metric | GO metric | BP | CC | MF | Joint* (pvalue<sup>+</sup>) |
| --- | --- | --- | --- | --- | --- |
| MHD | Resnik | 0.84194 | 0.77278 | 0.70436 | 0.86074 (2.93E-05) |
|  | Resnik<sub>RWC</sub> | 0.83743 | 0.75944 | 0.68886 | NA |
|  | AIC | 0.84042 | 0.76218 | 0.69590 | 0.85815 (5.93E-06) |
|  | GraSM | 0.76895 | 0.67114 | 0.64938 | NA |
|  | GraSM<sub>RWC</sub> | 0.76521 | 0.66967 | 0.63231 | NA |
|  | simDEF | 0.81356 | 0.72980 | 0.72644 | 0.84686 (1.80E-09) |
|  | W2vGO | 0.8246 | 0.79338 | 0.71915 | 0.86571 (4.66E-04) |
|  | InferSentGO | 0.83877 | 0.76026 | 0.75343 | 0.85368 (2.98E-07) |
|  | AicInferSentGO | 0.84830 | 0.78263 | 0.74489 | 0.86871 (2.05E-03) |
| BMA | Resnik | 0.85434 | 0.77871 | 0.70348 | 0.86766 (1.24E-03) |
|  | Resnik<sub>RWC</sub> | 0.84399 | 0.75626 | 0.68595 | NA |
|  | AIC | 0.85423 | 0.77628 | 0.69902 | 0.87854 (9.19E-02) |
|  | GraSM | 0.76895 | 0.67114 | 0.64938 | NA |
|  | GraSM<sub>RWC</sub> | 0.74201 | 0.64989 | 0.62591 | NA |
|  | simDEF | 0.83597 | 0.74432 | 0.72863 | 0.86958 (3.05E-03) |
|  | W2vGO | 0.84296 | **0.80035** | 0.71279 | 0.88170 (2.21E-01) |
|  | InferSentGO | 0.84600 | 0.76548 | **0.76081** | 0.87503 (2.85E-02) |
|  | AicInferSentGO | **0.85739** | 0.79011 | 0.74139 | **0.88987** |

To compare the performance of the metrics, we find the area under the curve (AUC) of the Receiver Operative Characteristic (ROC) curve. The real and random PPI networks serve as a basis to calculate the true positive and false negative rate, respectively. The AUC is computed by plotting the true positive versus false negative rates at different thresholds and estimating the area under this curve. AUC value goes from 0 to 1, with 1 being the best prediction power.

We judge the GO metrics based on their AUC values. From Fig. 2, because node-based methods adequately distinguish related BP terms from unrelated ones, the NLP methods’ improvement is best seen in the CC and MF ontologies (Table 2).

On average, each gene in this experiment is annotated by 20.66 GO terms, with the composition of 46.25% BP, 24.07% CC, and 29.68% MF terms. Because of these fractions, when using only the BP ontology to compare GO metrics, on average, we are using only 46.25% of the full description for a gene. The same argument can be made for using only the CC or MF ontology to compare GO metrics. For this reason, we conduct a joint analysis. Here, when comparing two genes, we use all the GO terms in their annotations, allowing for comparison of GO terms across different ontologies. This approach aligns with the observation that GO terms in different categories are connected (Fig. 1). Table 2 and 3 indeed show that the joint analyses yield the highest AUC for all GO metrics. We have two explanations for this outcome.

**Table 3:**
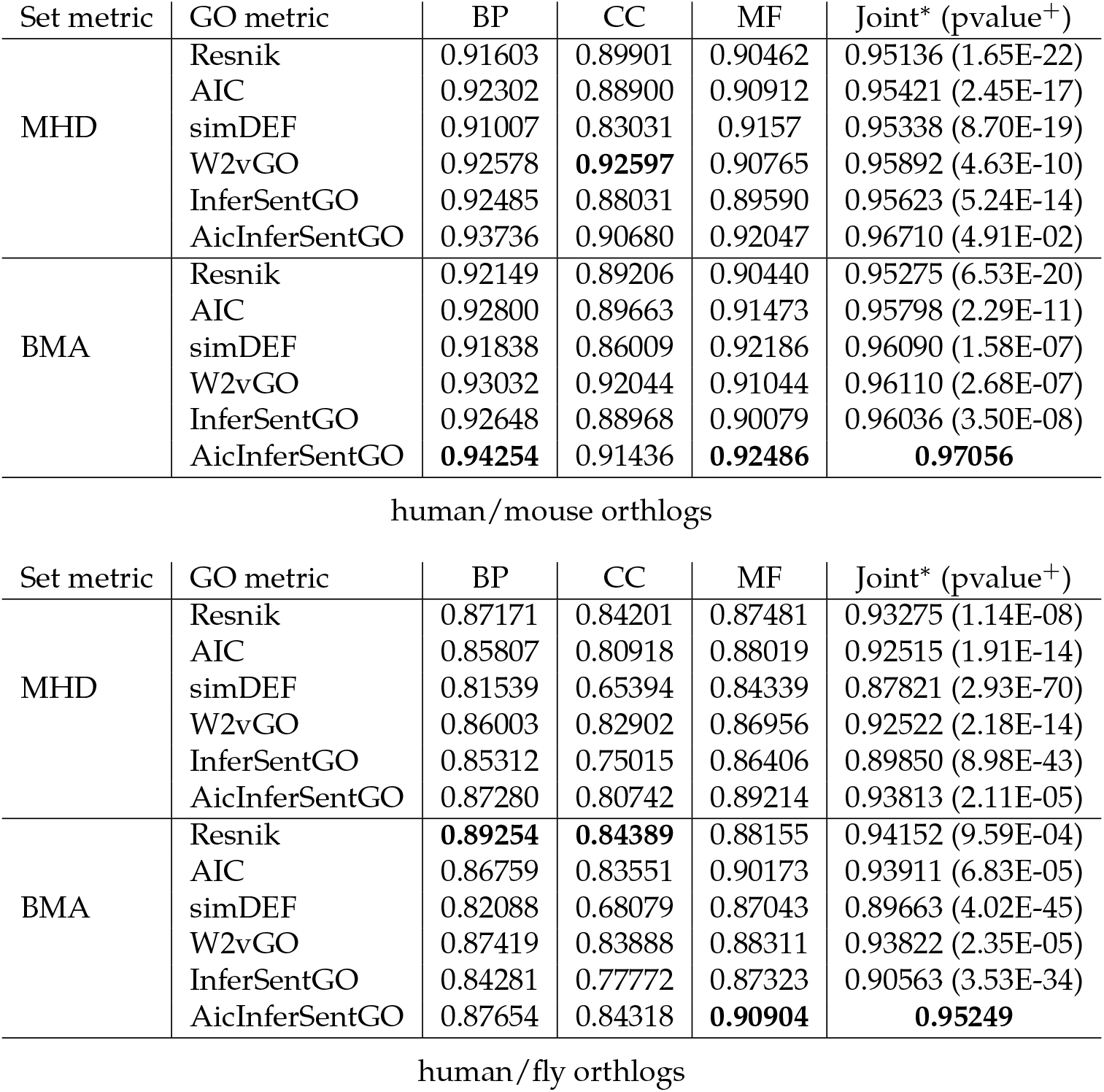
AUCs for classifying orthlogs. Bold font indicates the best value in each column. *Joint analysis: When comparing genes, we keep their entire GO annotations, effectively treating the BP, CC, MF ontologies as connected GO trees. +Pvalue of the Hanley-McNeil test to compare AUCs of other methods against BMA+AicInferSentGO.

First, intuitively, when using all BP, CC, and MF terms in the gene annotation, one can better understand the genes’ functionality. For example, when looking at the CC ontology terms alone, arguably proteins in the same part of the cell do not necessarily interact. However, when we consider not only the locations but also the biological and molecular events taking place, then we can accurately compare the two genes.

Second, empirically, in 46,967 randomly chosen child-parent pairs, we count 2060 pairs (4.38%) having terms in different ontologies. For example, a few terms having parents in different ontologies are GO:0009055, GO:0035514, GO:0102496, GO:1903198, and GO:1903934. The fraction 4.38%, despite being small, has a nontrivial repercussion.

This effect is especially true for Resnik and AIC whose key ideas rely on the number of common ancestors. For example, consider the term GO:0009055 in the MF ontology with its parent GO:0022900 in the BP category (Fig. 1). When treating the BP and MF trees separately, GO:0009055 and GO:0022900 have AIC score zero because they will not have any shared ancestors. When treating the trees jointly, the AIC score is 0.7197. Thus, we can better estimate the similarity between genes containing not only these terms but their descendant terms.

For reasons explained above, in this paper, we select the metric with the highest AUC in the joint analysis to be the best method. Here, Table 2 shows that BMA+AicInferSentGO does best. We note that BMA+W2vGO, despite not using information from the GO trees, works quite well on its own (2*^nd^* rank).

We use the Hanley-McNeil test to compare AUCs of the other methods against BMA+AicInferSentGO^[7]^. The p-values in column 6 of Table 2 show that BMA+AicInferSentGO is slightly better than BMA+AIC and BMA+W2vGO, and is statistically above the other approaches. We provide the ROC plots for the joint analyses at our GitHub.

Our joint analyses did not include Resnik_RWC_, GraSM, and GraSM_RWC_ because the software GOssTo does not allow the option to combine the three ontologies. In the experiment, we reimplemented Resnik and AIC ourselves, and changed the source code for simDEF (see GitHub).

### 3.2 Orthologs

Like in section 3.1, we remove electronically inferred GO terms from the gene annotation, and use genes with at least one GO term in each ontology. We test the following species: human/mouse and human/fly. For each pair, the positive set contains orthologs from the two species; whereas, the negative set contains randomly-matched genes. We set the sizes of the positive set and negative set to be equal. For human/mouse dataset, we have 10,235 pairs for each set; for the human/fly dataset, we have 4880 pairs for each set. We exclude Resnik_RWC_, GraSM, and GraSM_RWC_ in this experiment because they did not perform well for human protein network.

Like before, we consider the best GO metric to be the one with the highest AUC for the joint analysis. BMA+AicInferSentGO again does best in this experiment (Table 3). The Hanley-McNeil p-values for comparing AUCs indicate that BMA+AicInferSentGO is statistically above all other methods.

We also conduct the classification for mouse/fly orthologs. In the joint analysis, BMA+AicInferSentGO metric gives the highest AUC score 91.52% (full table not shown).

### 3.3 Yeast protein-protein interaction network

Pesaranghader <u>et al.</u>^[19]^ provided the yeast PPI data without electronically inferred GO terms. We keep only proteins annotated with at least one term in each ontology. The final dataset contains 3938 true interactions and 3938 random pairs. We consider the best GO metric to be the one with the highest AUC for the joint analysis. Here, Resnik outperforms the other GO metrics. We note that AicInferSentGO is better than both AIC and InferSentGO by themselves. This observation along with the previous results suggest that ensemble of node-based and NLP methods improves performance.

We hypothesize that text-based approaches (both ours and simDEF) do worst in part because PubMed data contains mostly papers on human biology. Evaluating the effect of training the model on different data sources is beyond the scope of this paper; we reserve this topic for future research.

## 4 Discussion

In our results, we do not aim to attain perfect classification; rather, we use the classification to rank the GO metrics. Other papers have used sequence similarity and co-expression data to evaluate GO metrics. However, sequence similarity has a stronger correlation to MF terms than BP and CC terms^[20]^. Also because of alternative splicing, similar sequences can produce proteins with different functionality^[25]^. Co-expression data works best with BP and CC ontology^[15,26]^, but genes are expressed non-uniformly across different tissues^[5]^. Depending on the data source, experiments using co-expression data can give highly varying outcomes.

The Word2vec has an extension Sentence2vec that converts a sentence into a vector^[10]^. Theoretically, one can convert GO definitions into vectors. However, our Word2vec result contains 986,615 words; so, the number of sentences in the training dataset is larger than this number. We encountered computer memory problem in training Sentence2vec on a 64GB RAM computer. Therefore, we opted for the InferSent model instead, training the model in 2 hours with GeForce GTX 1080 Ti 11GB graphic card.

Arguably, the *entailment* relation in InferSent does not necessarily equate to a perfect similarity measurement. For example, one can argue that every term in the BP ontology *entails* the root node *biological processes.* Moreover, the NLP approaches in this paper are yet to fully recognize chemical equations. An expression like 2*H*_2_ + *O*_2_ may not be seen as strictly equal to *2H*_2_*O* or the word *water.* For these reasons, we view NLP methods as ways to refine existing node-based GO metrics. In this paper, we have seen that InferSentGO improves the AIC scores. Moreover, InferSentGO does not need to be paired with AIC; it can work with any GO similarity metric that gives scores in the range [0, 1].

**Table 4:** AUCs for classifying yeast protein-protein interactions. Bold font indicates the best value in each column. *Joint analysis: When comparing genes, we keep their entire GO annotations, effectively treating the BP, CC, MF ontologies as connected GO trees. +Pvalue of the Hanley-McNeil test to compare AUCs of other methods against BMA+AicInferSentGO.

| Set metric | GO metric | BP | CC | MF | Joint* (pvalue<sup>+</sup>) |
| --- | --- | --- | --- | --- | --- |
| MHD | Resnik | 0.86739 | 0.83065 | 0.72235 | 0.89697 (1.16E-01) |
|  | Resnik<sub>RWC</sub> | 0.83743 | 0.75944 | 0.68886 | NA |
|  | AIC | 0.84422 | 0.78399 | 0.72323 | 0.85799 (7.03E-08) |
|  | GraSM | 0.70718 | 0.66114 | 0.66914 | NA |
|  | GraSM<sub>RWC</sub> | 0.71535 | 0.70451 | 0.56931 | NA |
|  | simDEF | 0.85859 | 0.72473 | **0.76989** | 0.87293 (4.38E-03) |
|  | W2vGO | 0.87192 | 0.80572 | 0.74163 | 0.88503 (4.96E-01) |
|  | InferSentGO | 0.87525 | 0.74827 | 0.74661 | 0.86655 (7.76E-05) |
|  | AicInferSentGO | 0.87059 | 0.77705 | 0.75379 | 0.87559 |
| BMA | Resnik | 0.88092 | **0.83759** | 0.72758 | **0.90592** (8.22E-04) |
|  | Resnik<sub>RWC</sub> | 0.84399 | 0.75626 | 0.68595 | NA |
|  | AIC | 0.86341 | 0.80490 | 0.73462 | 0.87426 (8.86E-03) |
|  | GraSM | 0.70454 | 0.64772 | 0.67181 | NA |
|  | GraSM<sub>RWC</sub> | 0.71198 | 0.69911 | 0.57286 | NA |
|  | simDEF | 0.83597 | 0.74432 | 0.72863 | 0.86958 (1.79E-01) |
|  | W2vGO | 0.87981 | 0.81583 | 0.74576 | 0.89118 (6.42E-01) |
|  | InferSentGO | 0.87832 | 0.76679 | 0.74788 | 0.87857 (6.40E-02) |
|  | AicInferSentGO | **0.88179** | 0.79532 | 0.76150 | 0.88870 |

We acknowledge that the literature contains many other methods for measuring GO terms’ semantic similarity which were not tested here. There has been much debate regarding which measure should be preferred over the others. However, no clear consensus has been reached^[29]^. In this paper, as a proof of concept, we show that NLP methods can work together with node-based models to achieve higher accuracy.

To our knowledge, this paper is the first to apply neural network based NLP techniques to compare the semantic meaning of GO terms. Our application suggests that there are great promises in developing NLP methods for this research area.

## Contributions

DD and JJL came up with the problem. DD and JJL decided on the data to be used. DD downloaded and cleaned the data. DD, JJL, WUA and KWC came up with the solution. DD and WUA coded the solution. DD wrote the paper. DD and JJL discussed the results. DD, JJL, WUA, KWC and EE read the paper.

## 5 Appendix

## 5.1 Resnik method

The most basic node-based method introduced by Resnik in 1999 relies on the information content (IC) of a GO term^[23]^. The IC of a GO term *t* is computed as IC(*t*) = − log(*p*(*t*)) where *p*(*t*) is the probability of observing a term *t* in the ontology. *p*(*t*) is computed as 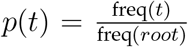. freq(*t*) is defined as the cumulative count of term *t* and its descendants, where 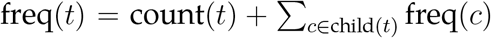. count(*t*) is the number of genes annotated with the term *t*, and child(*t*) are the children of *t*. Based on this definition, IC(*root*) = 0, and a node near the leaves has higher IC than nodes at upper levels. To compute a similarity score of the GO terms *a* and *b*, one finds the most informative common ancestor of these two terms.

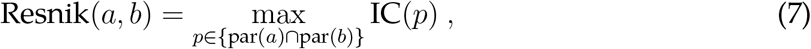

where par(*t*) denotes all the ancestors of term *t*. Resnik(*a, b*) ranges from 0 to ∞ because the probability *p*(*t*) ranges from 0 to 1.

In this model, the similarity score between a GO term *t* and itself is not 1. Second, when *a, b* have only the *root* as a common ancestor, then Resnik(*a, b*) = 0. This is problematic because leaf nodes are more informative than other types of nodes. Consider the example in Song <u>et al.</u>^[26]^. Here, the *root* is the only common ancestor of the pair *a, b* and the pair *c, d.* Next, suppose that *a, b* are leaf nodes, *c* is the parent of *a, d* is the parent of *b*, and *root* is the parent of both *c, d.* One would then expect that Resnik(*a, b*) < Resnik(*c*, *d*); however, one would obtain Resnik(*a, b*) = Resnik(*c*, *d*) = 0.

## 5.2 Aggregate Information Content (AIC) Method

The AIC method by Song <u>et al.</u>^[26]^ amends the two problems in the Resnik method. To encode the fact that leaf nodes are more informative, AIC defines a knowledge function of term *t* as *k*(*t*) = 1/IC(*t*) which is used to measure its semantic weight *sw*(*t*) = 1/(1 + exp(—*k*(*t*))). Here *sw*(*root*) = 1. Semantic value *sv*(*t*) of *t* is then defined as 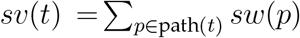. Function path(*t*) contains every ancestor of *t* and the term *t* itself. Usually, *sv*(*a*) *< sv*(*b*) when term *a* is nearer to the root than *b*. Song et al.^[26]^ define their similarity score of two GO terms *a, b* as

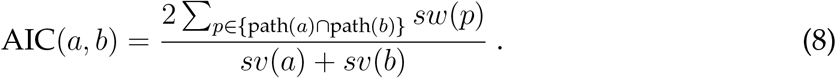

AIC(*a, b*) ranges from 0 to 1. In this model, AIC(*a*, *a*) = 1. When *a, b* have only the *root* as the common ancestor, then AIC(*a, b*) = 2/(*sv*(*a*) + *sv*(*b*)) which depends on where *a, b* are on the GO tree.

## 5.3 Graph-based Similarity Measure (GraSM)

GraSM is an extension of Resnik, and can be classified as an edge-based method. GraSM analyses more than just the most informative common ancestor of two GO terms, by looking at their disjunctive common ancestors^[3]^. For one GO term *a,* two of its ancestors are disjunctive if there are different paths from both ancestors to the GO term.

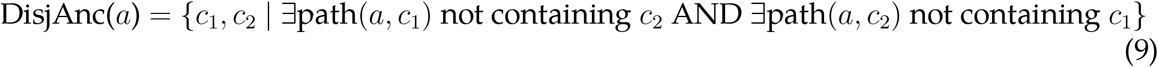

For two GO terms *a* and *b*, suppose the term *c*_1_ is in the union set *U*=DisjAnc(*a*)∪DisjAnc(*b*). *c*_1_ is a common disjunctive ancestor of *a, b* if for each common ancestor *c*_2_ of *a, b* where IC(*c*_1_) < IC(*c*_2_) we have both *c*_1_, *c*_2_ ∈ *U*. The similarity measurement for *a, b* is

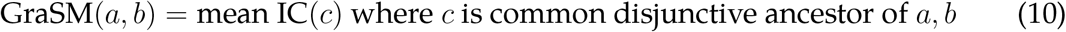

We use the software GOssTo to implement GraSM^[1]^.

## 5.4 Random Walk Contribution (RWC)

We briefly describe RWC’s key idea. Unlike many other methods which inspect the ancestors of two given GO terms, RWC is an edge-based approach that analyzes the shared children of two GO terms *a* and *b*^[29]^. In brief, in the RWC paradigm, GO terms with more common children are more deemed to be more similar.

Define *N_c_* as the number of genes annotated by term *c*. In RWC, the random walker moves from the parent node *p* to its direct child c with probability 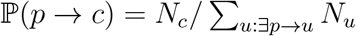. As the random walker moves for a very long time, we can denote the probability of ending at a node *i* from *a* to be 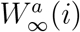. Let *L* be the set of all leaf nodes in the GO tree. The RWC for two terms *a* and *b* is

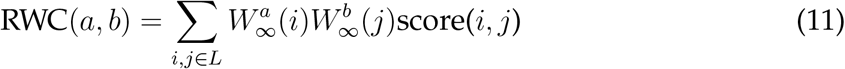

score(*i*, *j*) is a generic place holder. For example, score(*i*, *j*) can be Resnik(*i*, *j*). Yang <u>et al.</u>^[29]^ uses their RWC to improve score(*a, b*) by taking the average

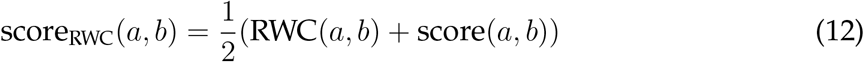

In this paper, we use the software GOssTo to implement RWC with score(*a, b*) being Resnik and GraSM^[1]^. Currently, GOssTo is unable to take any generic score function as its argument.

## Footnotes

1 The original authors did not name their method; hence, we use their GitHub name. The NLP community refers to it as bi-directional long short-term memory with max-pooling.

2 github.com/datduong/NLPMethods2CompareGOterms

3 The user specifies *N*; in practice, people often set *N* = 300.

